# One Molecule, Multiple Histories: Comparative Phylogenomics of Glutamate Metabolism, Transport and Receptor Families Across 89 Eukaryotic Proteomes

**DOI:** 10.64898/2026.09.05.749567

**Authors:** Ankit Thakur, Mahesh Kulharia

**Affiliations:** Centre for Computational Biology and Bioinformatics, Central University of Himachal Pradesh, Dharamshala, District Kangra, Himachal Pradesh - 176215, INDIA

**Keywords:** Glutamatergic evolution, comparative phylogenomics, human-reference-anchored homology, Molecular evolution, Neurotransmitter evolution, Gene-tree/species-tree reconciliation

## Abstract

Glutamate links cellular metabolism with excitatory neurotransmission, yet the evolutionary histories of its metabolic enzymes, transporters, and receptor paralogues have not been compared within a common human-reference-anchored phylogenomic framework. Here, we traced human UniProt/Swiss-Prot-associated protein lineages across 89 eukaryotic proteomes using homology searches, sequence and domain filtering, maximum-likelihood phylogenies assessed with transfer bootstrap expectation (TBE) and ultrafast bootstrap (UFBoot), sequence-stability analyses, and GeneRax gene-tree/species-tree reconciliation under a duplication–loss model. Metabolic enzyme homologues showed the broadest distributions: GLUD, GOT, and GPT were recovered across multiple eukaryotic supergroups, whereas GLS-associated sequences were detected in metazoans and the ichthyosporean *Capsaspora owczarzaki*. SLC17 and SLC1 transporter homologues were recovered only from sampled metazoans. Receptor distributions varied among families: GRM and GRIN sequences were detected across sampled cnidarians and bilaterians, GRIA and GRIK across bilaterian groups, and GRID only in seven sampled vertebrate species. Reconciliation mapped numerous paralogue-associated duplications to early vertebrate, gnathostome, and osteichthyan ancestral nodes, alongside independent lineage-specific expansions. Although many local phylogenetic relationships were supported, deeper relationships in some transporter and receptor families remained uncertain, with TBE and UFBoot providing different support values. These findings characterize the human-associated glutamatergic repertoire as an evolutionary mosaic of gene families with distinct taxonomic distributions and duplication histories. Because homology searches were anchored on human reference proteins, the inferred distributions represent detectable lineages in the sampled dataset rather than definitive evidence of gene-family origins or absence.

## Introduction

Glutamate links cellular metabolism to intercellular signaling: it is an abundant metabolite at the intersection of carbon and nitrogen metabolism and the primary excitatory transmitter in the mammalian brain. Glutamate-mediated signaling is also found beyond synapses, including in early-branching animals and unicellular eukaryotes [1], [2]. At synapses, vesicular glutamate transporters (VGLUTs) load glutamate into synaptic vesicles; after release, glutamate activates ionotropic and metabotropic receptors, while excitatory amino-acid transporters (EAATs) take it up from the extracellular space [3], [4]. Glutaminase forms glutamate from glutamine, while glutamate dehydrogenase and aminotransferases mediate glutamate metabolism, linking glutamate turnover to carbon and nitrogen metabolism [5], [6]. Together, vesicular loading, receptor activation, uptake, and metabolism couple transmitter availability and synaptic signaling with extracellular glutamate homeostasis; vesicular and plasma-membrane transport are distinct molecular systems [7], [8].

Comparative genomics has challenged a vertebrate-centered view of glutamatergic signaling. Ionotropic glutamate-receptor-like genes have been reported in plants; an early phylogenetic analysis placed plant GLRs within the broader iGluR lineage and proposed that plant–animal receptor divergence preceded the diversification of animal iGluR subtypes [9]. Subsequent metazoan-wide analyses found that vertebrate-based classifications do not capture the full diversity of animal iGluRs and reported lineage-specific evolutionary histories [10]. Metabotropic glutamate-receptor GPCRs also have deep roots within animals: comparative surveys identified GRM-family receptors in invertebrates, including *Caenorhabditis elegans* and *Ciona intestinalis*, as well as vertebrates [11]. Together, these studies show that human receptor families sit within a broader and more diverse evolutionary context; however, these analyses do not resolve the histories of every human-associated paralogue or establish a single origin for the complete glutamatergic system [9], [10], [11].

Transporter studies reveal family-specific evolutionary histories. Phylogenetic analysis of SLC17 identified four major clades that were already present before insect divergence; one clade was not found in vertebrates [12]. In SLC1, a survey of more than 300 genes across more than 40 vertebrate species combined phylogeny and microsynteny, identifying lineage-specific subfamilies and evidence that therian mammals lost several genes retained in other vertebrate lineages; some additional ray-finned fish copies were attributed to the third-round, teleost- specific genome duplication [13]. For the GRIN2 receptor-subunit family, phylogeny, synteny and paralogon analyses supported the origin of the four-gene vertebrate set through two rounds of genome duplication, with additional fish copies associated with the third-round fish-specific duplication [14]. Together, these cases show that genome duplication explains some family expansions, while lineage-specific retention and loss also contribute to present-day repertoires; they do not justify attributing every paralogue or family history to whole-genome duplication [13], [14].

Metabolic-enzyme histories likewise caution against a single family-wide model. GLUD2 arose by retroposition from GLUD1 in the hominoid ancestor less than 23 million years ago, and amino-acid changes associated with its brain-specific properties accumulated under positive selection after duplication [15]. This example distinguishes the deeper history of a gene family from the more recent origin and subsequent evolution of a particular paralogue. However, these studies address individual protein families or subfamilies, leaving unresolved how the evolutionary histories of human-reference-associated metabolic enzymes, transporters, and receptor paralogues compare within a common phylogenomic framework. Moreover, an apparent absence in an annotated genome may reflect sequencing or gene-prediction error rather than a true gene loss, so absence from sampled proteomes should be treated cautiously [16].

A comparative framework spanning these functional classes can assess how far human-reference-associated sequences can be traced across eukaryotes, whether supported gene-tree relationships align with the species tree, and how uncertainty in sequence placement affects inferred duplication and loss. In this study, human UniProt/Swiss-Prot proteins from glutamate-metabolic enzymes, vesicular and plasma-membrane transporters, and ionotropic and metabotropic receptors were used to identify homologues across 89 eukaryotic species. UniProt provides curated protein sequences and functional annotations, although Swiss-Prot inclusion does not itself establish orthology or evolutionary history [17]. Maximum-likelihood gene trees were assessed using Transfer Bootstrap Expectation and ultrafast bootstrap support [18]; leaf-stability and t-index assessments were used to evaluate sensitivity to taxon placement. GeneRax was used for species-tree-aware gene-tree inference and reconciliation under gene duplication, and loss [19]. The aim is an integrated, uncertainty-aware comparison of the inferred histories of these human-associated sequences, not a claim to establish the universal origin of glutamate signalling or definitive gene absence where homologues were not detected.

## Results

### Evolution of glutamate-metabolic enzymes combines ancient origins with lineage-specific duplications

Glutamate homeostasis involves two mitochondrial enzyme families: glutaminases (GLS and GLS2), which hydrolyze glutamine to glutamate and ammonia, and glutamate dehydrogenase (GLUD1 and GLUD2), which reversibly interconvert glutamate and α-ketoglutarate with the release or assimilation of ammonia. To reconstruct the evolutionary histories of these families, we inferred maximum-likelihood (ML) phylogenies, assessed nodal support using ultrafast bootstrap (UFB) and transfer bootstrap expectation (TBE), evaluated sequence-tip stability using the leaf stability index (LSI) and t-index, and reconciled the inferred gene trees with the species tree using GeneRax.

First, we investigated the glutaminase family, represented in humans by GLS (UniProt O94925; kidney/brain-type) and GLS2 (UniProt Q9UI32; liver-type). Our phylogenetic trees recovered homologs across opisthokonts (*Capsaspora owczarzaki*), non-bilaterians (*Bolinopsis microptera, Amphimedon queenslandica, Trichoplax sp. H2*, and anthozoan cnidarians), and bilaterians (Fig. 1A; Online Resource J). Eight sequence tips were flagged as unstable by the LSI and t-index analyses and excluded before reconciliation. Their sequence identifiers and taxonomic assignments are provided in Online Resource K. In the phylogenetic trees, jawed vertebrate (gnathostome) glutaminases formed a monophyletic group (TBE=0.95, UFB=65) sister to lampreys (TBE=0.98, UFB=100), and segregated into two well-supported clades: the GLS clade (TBE=1.0, UFB=98), containing human GLS within a monophyletic tetrapod cluster (TBE=0.98, UFB=82), and the GLS2 clade (TBE=0.95, UFB=88), containing human GLS2 (TBE=0.99, UFB=100) (Fig. 1a; Online Resource J). After excluding unstable sequence tips, GeneRax placed the deepest represented glutaminase-family node at an ancestral node encompassing the sampled metazoan and ichthyosporean lineages. This represents the deepest node recovered in the sampled dataset rather than establishing a definitive origin at the Filozoa stem. The reconciliation inferred an independent expansion in the ctenophore *Bolinopsis microptera* (11 duplications), as well as secondary losses in Medusozoa and *Drosophila melanogaster*. The lineages corresponding to human GLS and GLS2 were traced to a duplication in the jawed-vertebrate ancestor, followed by inferred lineage-specific losses of GLS in *Monodelphis domestica* and GLS2 in *Gallus gallus* (Fig. 1b; Online Resource M(b)).

**Fig. 1.**
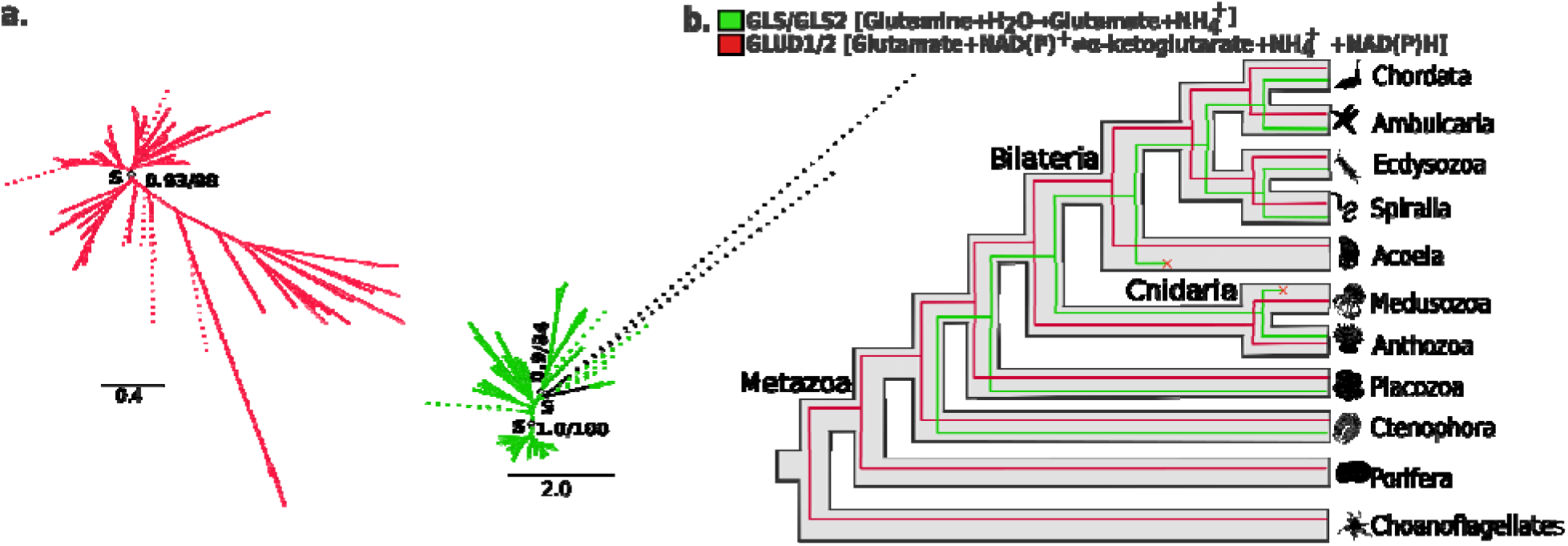
Phylogenetic relationships and gene-tree-species-tree reconciliation of glutaminases (GLS and GLS2) and glutamate dehydrogenases (GLUD1 and GLUD2). (a) Transfer bootstrap expectation trees showing the evolutionary relationships of the GLS and GLUD protein families. Nodal support values are shown as transfer bootstrap expectation (TBE; decimal values) and ultrafast bootstrap (UFB; whole-number percentages) for selected nodes. Dashed branches indicate sequences identified as unstable by the t-index or leaf stability index (LSI) analyses. (b) Simplified, combined representations of the corresponding gene-tree-species-tree reconciliations inferred using GeneRax. Asterisks (✰) indicate speciation (S) or duplication (D) events, with TBE and UFB support values provided where applicable. Species silhouettes were obtained from PhyloPi (https://www.phylopic.org/). Complete annotated phylogenetic trees and nodal support values are provided in Online Resource J, and detailed GeneRax reconciliation outputs are provided in Online Resource M(b).

Next, we examined the glutamate dehydrogenase family, represented in humans by GLUD1 (UniProt P00367) and GLUD2 (UniProt P49448). Our phylogenetic analyses indicated that GLUD enzymes are conserved across eukaryotes, spanning plants, amoebozoans, fungi, choanoflagellates, non-bilaterians, and bilaterians. Ten sequence tips were flagged as unstable by the LSI and t-index analyses and excluded before reconciliation; their identifiers and taxonomic assignments are provided in Online Resource K. Both TBE and UFB trees recovered the monophyly of chordate (TBE=0.92, UFB=92), vertebrate (TBE=0.99, UFB=100), gnathostome (TBE=0.95, UFB=93), and mammalian (TBE=0.93, UFB=80) GLUDs (Fig. 1a; Online Resource J). Human GLUD1 and GLUD2 were recovered in a human-associated clade near mouse Glud1 in the sampled phylogeny (Fig. 1a; Online Resource J). GeneRax otherwise indicated broadly species-tree-concordant evolution across Metazoa, with additional lineage-specific duplications, including glud1a and glud1b in *Danio rerio* (Fig. 1b; Online Resource M(b)). However, the placement of the GLUD1/GLUD2 duplication on the Homo sapiens branch in the present analysis should not be interpreted as establishing a human-specific origin, because comparative evidence places the origin of GLUD2 by retroposition of GLUD1 in the hominoid ancestor.

### Evolutionary History of Glutamate Catabolic Enzymes

Tight regulation of synaptic and intracellular glutamate levels is essential to prevent excitotoxicity and maintain metabolic homeostasis. In humans, the transamination-mediated catabolism and cycling of glutamate are catalyzed by two pyridoxal phosphate-dependent enzyme families: aspartate aminotransferases (GOT / AAT) and alanine aminotransferases (GPT / ALAT). Following the same phylogenomic and reconciliation pipeline, we examined the evolutionary origin and duplication patterns of the human catabolic repertoire.

#### Aspartate Aminotransferases (GOT1, GOT1L1, and GOT2)

Aspartate aminotransferases catalyze the reversible transfer of an α-amino group from aspartate to α-ketoglutarate, producing oxaloacetate and glutamate. Human GOT1 (UniProt P17174) wa used as the initial seed query for the aspartate aminotransferase analysis. Downstream sequence recovery and family-level analyses identified the human GOT1, putative GOT1L1 (UniProt Q8NHS2), and mitochondrial GOT2 (UniProt P00505) proteins for inclusion in the comparative analysis. Both TBE and UFB phylogenetic reconstructions demonstrated that the GOT family i conserved across all major eukaryotic supergroups (Fig. 2a; Online Resource J), resolving two ancient clades corresponding to the mitochondrial GOT2 and cytoplasmic GOT1 lineages. Four mitochondrial-like sequences from the sponge *A. queenslandica* were flagged as unstable: they grouped with *D. melanogaster* GOT2 in the TBE tree but with amoebozoans in the UFB tree. These sequences were excluded before reconciliation (Online Resource K).

**Fig. 2.**
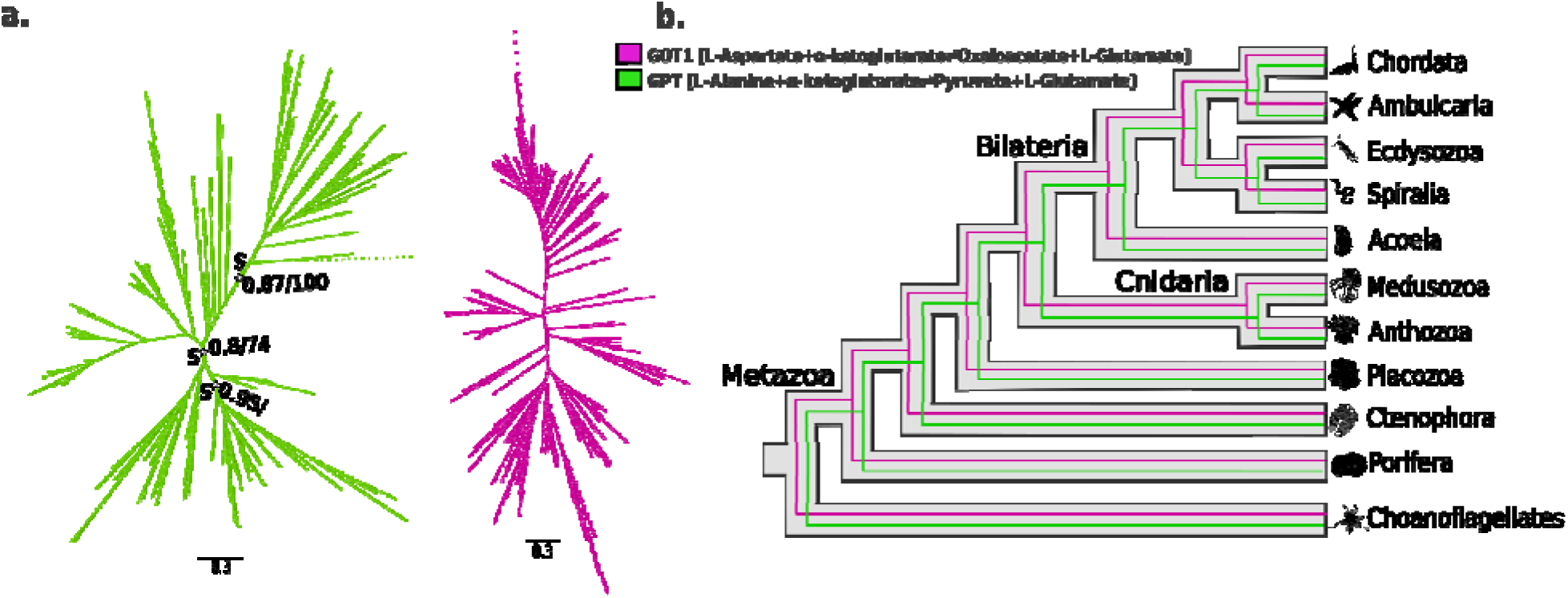
Phylogenetic relationships and gene-tree-species-tree reconciliation of aspartate aminotransferases (GOT1, GOT1L1, and GOT2) and alanine aminotransferases (GPT and GPT2). (a) Transfer bootstrap expectation (TBE) phylogenetic trees depicting the evolutionary relationships of the GOT and GPT protein families, including GOT1, GOT1L1, GOT2, GPT, and GPT2. Nodal support value are shown as transfer bootstrap expectation (TBE; decimal values) and ultrafast bootstrap (UFB; whole-number percentages) for selected nodes. Dashed branches indicate sequences identified as unstable b the t-index or leaf stability index (LSI) analyses. (b) Simplified, combined representations of the corresponding gene-tree-species-tree reconciliations inferred using GeneRax. Asterisks (✰) indicate speciation (S) or duplication (D) events, with TBE and UFB support values provided where applicable. Species silhouettes were obtained from PhyloPic (https://www.phylopic.org/). Complete annotated phylogenetic trees and nodal support values are provided in Online Resource J, and detailed GeneRax reconciliation outputs are provided in Online Resource M(b).

Within the mitochondrial GOT2 clade, vertebrate sequences formed a strongly supported group (TBE = 1.00; UFB = 100) that included lampreys and jawed vertebrates (TBE = 0.89; UFB = 75). The mammalian GOT2 clade, including the human protein, was also strongly supported (TBE = 1.00; UFB = 100). In the cytoplasmic GOT1 clade, chordate homologues formed a supported monophyletic group (TBE = 0.96; UFB = 85). Tunicate and lamprey sequences were positioned outside the jawed-vertebrate grouping, whose support was discordant between TBE (0.95) and UFB (45). Within jawed vertebrates, sequences bifurcated into two paralogous subclades: the canonical GOT1 clade (TBE=0.96, UFB=85), containing human GOT1 (P17174) within a monophyletic tetrapod (TBE=0.81, UFB=81) and mammalian (TBE=0.92, UFB=97) cluster; and the GOT1L1 (AATC2) clade (TBE=1.0, UFB=100), comprising zebrafish, coelacanth, lizard (Anolis carolinensis), and mammalian GOT1L1 (including human GOT1L1, Q8NHS2; TBE=1.0, UFB=100) (Fig. 2a; Online Resource J). GeneRax reconciliation inferred an ancient divergence between the cytoplasmic GOT1 and mitochondrial GOT2 lineages, predating the emergence of Metazoa. GOT2 was retained largely as a single-copy gene across vertebrates, apart from inferred lineage-specific duplications in *Danio rerio* and *Gallus gallus*. GeneRax placed the GOT1/GOT1L1 duplication at the osteichthyan ancestral node represented by mouse and zebrafish, followed by inferred losses of GOT1L1 in *Xenopus tropicalis, Gallus gallus*, and *Takifugu rubripes* (Fig. 2b; Online Resource M(b)).

#### Alanine Aminotransferases (GPT and GPT2)

Alanine aminotransferases catalyze the reversible transamination between alanine and α-ketoglutarate, producing pyruvate and glutamate. Human GPT (UniProt P24298) was used as the initial seed query for the alanine aminotransferase analysis. The downstream analysis also recovered the human GPT2 protein (UniProt Q8TD30), which was included in the comparative phylogenetic analysis. Phylogenetic analyses (TBE and UFB) revealed that the GPT orthogroup is widely distributed across eukaryotes, including plants, fungi, amoebozoans, non-bilaterians, and bilaterians. Only a single partial sequence from the telonemid *Telonema subtile* was flagged as unstable by the LSI and t-index analyses and removed before reconciliation (Online Resource K).

The inferred GPT phylogeny recovered a large metazoan assemblage. Chordate sequences, including tunicate and vertebrate homologues, clustered with TBE = 0.95 and UFBoot = 97, while vertebrate homologues formed a monophyletic clade with the same reported support values. Bony vertebrate sequences split into two major paralogous groups corresponding to the human enzymes:

The GPT (ALAT1) Clade: The clade containing human GPT (P24298) was strongly supported (TBE=0.92, UFB=95), pairing a teleost-duplicated subclade (TBE=0.98, UFB=100) with a sarcopterygian (coelacanth + tetrapod) clade (TBE=0.90, UFB=60), within which tetrapod (TBE=0.95, UFB=93) and mammalian (TBE=1.0, UFB=100) GPT sequences were monophyletic.

The GPT2 (ALAT2) Clade: The sister clade (TBE=1.0, UFB=100) encompassed teleost, cartilaginous fish, coelacanth, and tetrapod GPT2 sequences (TBE=0.98, UFB=97), recovering mammalian GPT2 (including human GPT2, Q8TD30) as a robust monophyletic unit (TBE=1.0, UFB=100). A third, more basal gnathostome/cyclostome paralogous cluster (TBE=0.68, UFB=50) was retained only in fishes, coelacanth, and lampreys (Fig. 2a; Online Resource J). GeneRax placed the deepest represented GPT-family node at the common ancestor represented by mouse and Thecamonas trahens in the sampled species tree. This placement identifies the deepest node recovered in the analysis but does not, by itself, establish an origin at the root of Eukaryota. Within Metazoa, independent lineage-specific expansions were inferred in cnidarians (e.g., *Hydra vulgaris, Pocillopora verrucosa*) and arachnids (*Argiope bruennichi*). GeneRax mapped four duplication events to the ancestral node represented by mouse and zebrafish, corresponding to the osteichthyan ancestor. It also inferred the loss of a GPT-associated copy in *Gallus gallus* and *Callorhinchus milii*, as well as the loss of the third basal paralogue in the tetrapod ancestor. However, the reported loss in *Callorhinchus milii* requires verification against the reconciled tree before the precise timing of GPT/GPT2 divergence can be established (Fig. 2b; Online Resource M(b)).

### Evolutionary History of Glutamate Transporters

In humans, glutamate-mediated neurotransmission depends on two major transporter families: vesicular glutamate transporters (VGLUTs), which package glutamate into synaptic vesicles, and excitatory amino acid transporters (EAATs), which mediate glutamate uptake and contribute to the regulation of extracellular glutamate levels. Using human UniProt/Swiss-Prot proteins as query sequences, we investigated the evolutionary origins and duplication patterns of both transporter families across the 89-species eukaryotic dataset.

#### Vesicular Glutamate Transporters (SLC17A6, SLC17A7, and SLC17A8)

First, we investigated the vesicular glutamate transporter family, represented in humans by VGLUT1 (SLC17A7; UniProt Q9P2U7), VGLUT2 (SLC17A6; UniProt Q9P2U8), and VGLUT3 (SLC17A8; UniProt Q8NDX2). Our orthogroup analysis recovered 140 sequences from diverse metazoan lineages, including cnidarians, the placozoan *Trichoplax sp. H2*, and bilaterians. No homologues were detected in the sampled non-metazoan proteomes. Both TBE and UFB phylogenies recovered a strongly supported assemblage containing the three human reference proteins (TBE = 1.00; UFB = 100). Within this assemblage, the SLC17A7- and SLC17A6-associated lineages grouped together with moderate-to-strong support (TBE = 0.83; UFB = 84). The human VGLUT1 sequence clustered with mouse VGLUT1 (TBE = 1.00; UFB = 100), while human VGLUT2 clustered with mouse VGLUT2 (TBE = 0.94; UFB = 86) and human VGLUT3 with mouse VGLUT3 (TBE = 0.97; UFB = 98). These relationships support the separation of the vertebrate-associated VGLUT paralogue groups, although the broader family also contains sequences from diverse non-bilaterian and bilaterian lineages (Fig. 3a; Online Resource J). No SLC17A sequences were flagged as unstable by the LSI or t-index analyses, and all recovered sequences were retained for reconciliation.

**Fig. 3.**
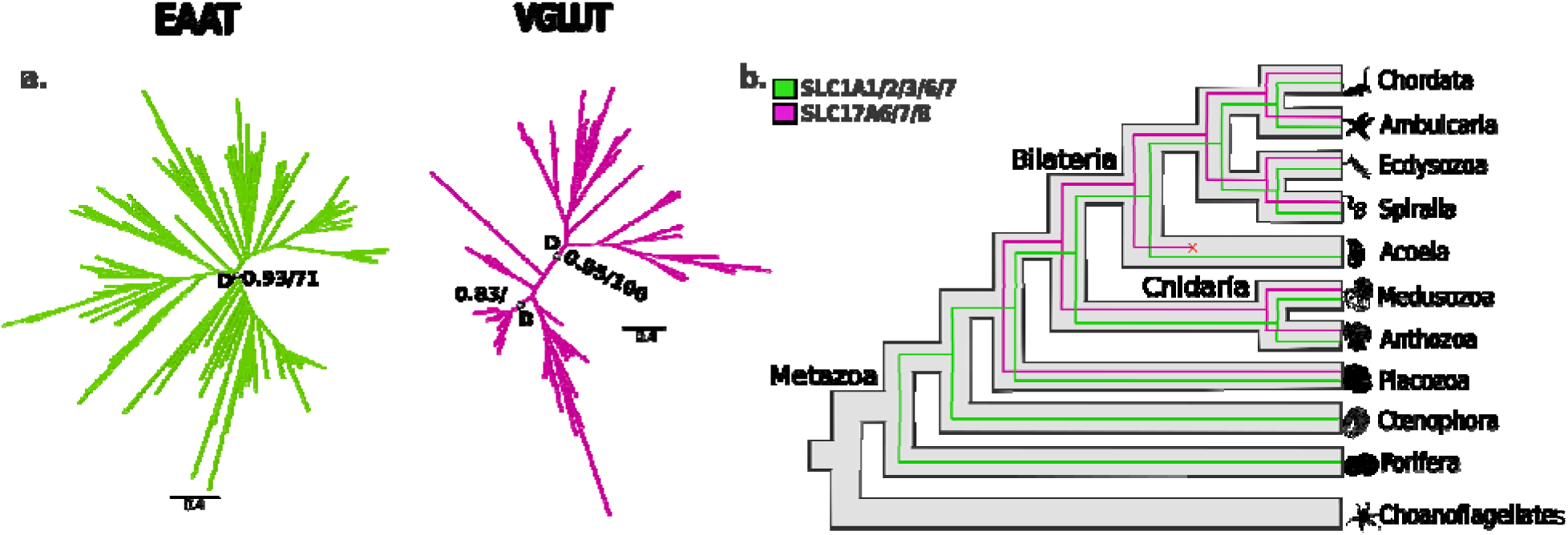
Phylogenetic relationships and gene-tree-species-tree reconciliation of vesicular glutamate transporters (VGLUTs; SLC17A6, SLC17A7, and SLC17A8) and excitatory amino acid transporters (EAATs; SLC1A1, SLC1A2, SLC1A3, SLC1A6, and SLC1A7). (a) Transfer bootstrap expectation (TBE) phylogenetic trees depicting the evolutionary relationships of the SLC17A and SLC1A protein families, including SLC17A6, SLC17A7, SLC17A8, SLC1A1, SLC1A2, SLC1A3, SLC1A6, and SLC1A7. Nodal support values are shown as transfer bootstrap expectation (TBE; decimal values) and ultrafast bootstrap (UFB; whole-number percentages) for selected nodes. (b) Simplified, combined representations of the corresponding gene-tree-species-tree reconciliations inferred using GeneRax. Asterisks (✰) indicate speciation (S) or duplication (D) events, with TBE and UFB support values provided where applicable. Species silhouettes were obtained from PhyloPic (https://www.phylopic.org/). Complete annotated phylogenetic trees and nodal support values are provided in Online Resource J, and detailed GeneRax reconciliation outputs are provided in Online Resource M(b).

GeneRax reconciliation inferred a duplication-rich evolutionary history, with events mapped to several ancestral lineages, including the common ancestors represented by mouse and Acropora muricata, mouse and Strongylocentrotus purpuratus, and mouse and Petromyzon marinus. Additional lineage-specific duplications were inferred within cnidarians, ecdysozoa, and teleosts. Crucially, GeneRax mapped the duplications separating the SLC17A8-associated lineage from the SLC17A6/SLC17A7-associated lineages to the gnathostome ancestor. This pattern supports diversification of the three lineages represented by modern human VGLUT paralogues during early jawed-vertebrate evolution, against a background of older and independent duplications elsewhere in the family (Fig. 3b; Online Resource M(b)).

#### Excitatory Amino Acid Transporters (SLC1A1, SLC1A2, SLC1A3, SLC1A6, and SLC1A7)

Next, we examined the evolutionary history of the excitatory amino acid transporter family using the five human reference proteins EAAT1 (SLC1A3, P43003), EAAT2 (SLC1A2, P43004), EAAT3 (SLC1A1, P43005), EAAT4 (SLC1A6, P48664), and EAAT5 (SLC1A7, O00341). Our orthogroup analysis recovered 286 sequences distributed across diverse metazoan lineages, including the sponge Sycon ciliatum, the ctenophore Bolinopsis microptera, cnidarians, the placozoan *Trichoplax sp. H2*, and multiple protostomes and deuterostome bilaterians. Despite sampling across the broader eukaryotic dataset, no non-metazoan sequences were recovered in this orthogroup. The inferred phylogenetic topologies were broadly concordant, but TBE and UFBoot assigned different levels of support to some deeper relationships within the EAAT family. In particular, the broad grouping containing the human SLC1A1 and SLC1A2 reference sequences received support of TBE = 0.98 in the TBE tree, whereas the corresponding grouping was weakly supported in the UFB tree (UFB = 22). Similarly, the node containing the human SLC1A3 and SLC1A6 reference sequences received TBE = 0.90 and UFB = 67. These differences indicate that, although several local relationships within the EAAT family were supported, its deeper paralogue relationships were not consistently resolved by both phylogenetic reconstructions. No sequences in either transporter orthogroup were flagged as unstable by the LSI or t-index analyses; all recovered sequences were retained for reconciliation (Fig. 3a; Online Resource J).

GeneRax placed the deepest represented EAAT-family node at the common ancestor of mouse and the sponge *A. queenslandica*. Additional duplication events were mapped to ancestral nodes represented by the common ancestors of cnidarians, mouse and amphioxus (*Branchiostoma belcheri*), mouse and lamprey, and mouse and elephant shark (*Callorhinchus milii*), indicating successive duplication events throughout metazoan, chordate, and vertebrate evolution. In particular, the divergence of the lineages containing the human SLC1A3 and SLC1A6 reference sequences was mapped to the gnathostome ancestral node. Further duplication events were assigned to sarcopterygian and tetrapod ancestral nodes (Fig. 3b; Online Resource M(b)). Overall, the EAAT-family tree supports a metazoan origin followed by repeated duplications and lineage-specific losses. Discordance between TBE and UFBoot support values, however, prevents a confident reconstruction of the sequence of events that produced all five human paralogues (Fig. 3a; Online Resource J).

### Evolutionary History of Glutamate Receptors

Glutamate signaling is mediated by ionotropic AMPA-type (GRIA), kainate-type (GRIK), delta-type (GRID), and NMDA-type (GRIN) receptors, as well as metabotropic glutamate receptors (GRM). To reconstruct the evolutionary histories of these families, we inferred maximum-likelihood phylogenies using human UniProt/Swiss-Prot reference proteins, assessed nodal support using transfer bootstrap expectation (TBE) and ultrafast bootstrap (UFB), evaluated sequence-tip stability using LSI and t-index analyses, and reconciled the resulting gene trees with the species tree using GeneRax. The analyses covered all five receptor families. No sequences in the GRIA, GRIK, GRID, GRM, or GRIN orthogroups were flagged as unstable by the LSI or t-index analyses; all recovered sequences were retained for reconciliation.

#### AMPA-Type Glutamate Receptors (GRIA1-GRIA4)

First, we investigated the AMPA-type ionotropic glutamate receptor family using the four human reference proteins GRIA1 (P42261), GRIA2 (P42262), GRIA3 (P42263), and GRIA4 (P48058). Our orthogroup analysis recovered 64 sequences from 27 species, spanning diverse bilaterian lineages, including chordates, echinoderms, ecdysozoa, molluscs, annelids, and the rotifer Adineta vaga. Both TBE and UFB phylogenies recovered vertebrate-associated groups containing sequences assigned to the four GRIA paralogues. The group containing human GRIA1 and its vertebrate-associated homologues was strongly supported (TBE = 1.00; UFB = 100). A broader vertebrate grouping containing sequences assigned to GRIA1-GRIA4 was also strongly supported (TBE = 1.00; UFB = 100). Within this assemblage, a grouping containing GRIA2-, GRIA3-, and GRIA4-associated sequences received TBE = 0.94 and UFB = 93, whereas a broader relationship involving additional chordate and echinoderm sequences received lower support (TBE = 0.92; UFB = 63). Thus, vertebrate-associated GRIA lineages were consistently recovered, whereas deeper relationships among the paralogue-associated groups remained less well resolved (Fig. 4a; Online Resource J).

**Fig. 4.**
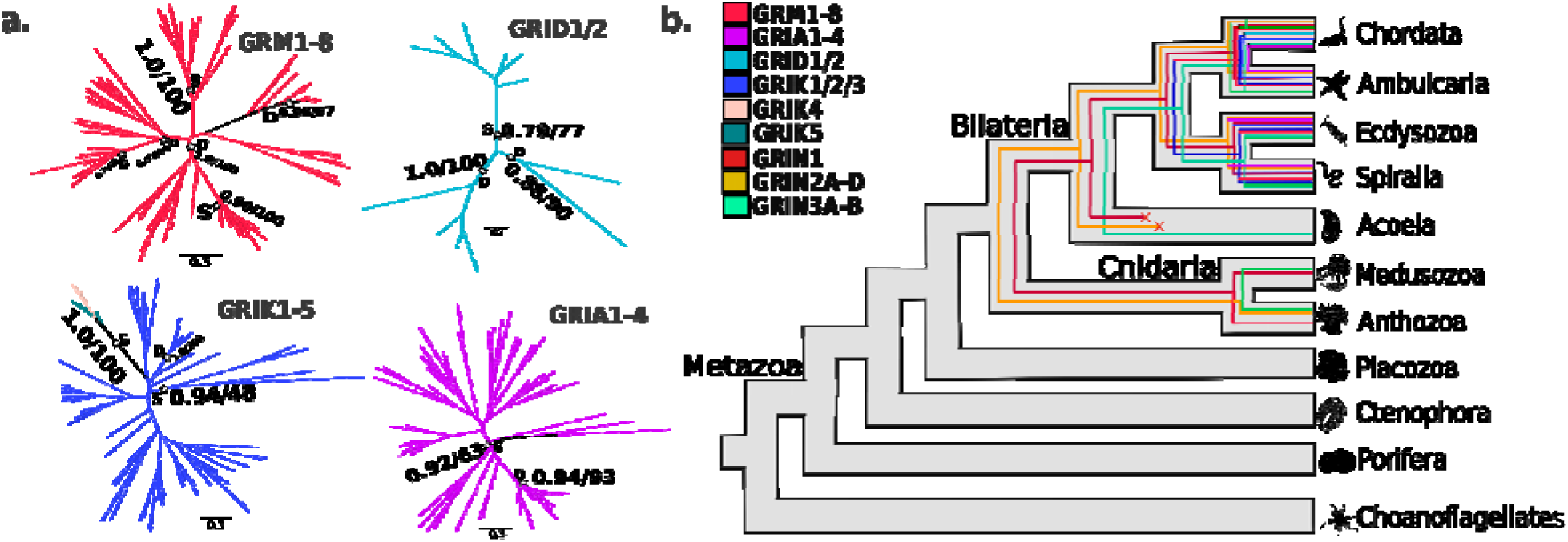
Phylogenetic relationships and gene-tree-species-tree reconciliation of ionotropic and metabotropic glutamate receptors. (a) Transfer bootstrap expectation (TBE) phylogenetic trees depicting the evolutionary relationships of the AMPA-type (GRIA), kainate-type (GRIK), and delta-type (GRID) ionotropic glutamate receptor families, together with the metabotropic glutamate receptor (GRM) family. Nodal support values are shown a transfer bootstrap expectation (TBE; decimal values) and ultrafast bootstrap (UFB; whole-number percentages) for selected nodes. (b) Simplified, combined representations of the corresponding gene-tree-species-tree reconciliations inferred using GeneRax. Asterisks (✰) indicate speciation (S) or duplication (D) events, with TBE and UFB support values provided where applicable. Species silhouette were obtained from PhyloPic (https://www.phylopic.org/). Complete annotated trees and nodal support values for all receptor families, including the separate GRIN phylogeny, are provided in Online Resource J; detailed GeneRa reconciliation outputs are provided in Online Resource M(b).

GeneRax reconciliation inferred 31 duplication events, 32 speciation events, and 28 speciation-loss events, no standalone-loss events. The root of the reconciled tree was mapped to the node represented by the common ancestor of mouse and *Symsagittifera roscoffensis*, consistent with a bilaterian ancestral history. Duplication events were distributed across multiple lineages, including three events at the ancestral node represented by mouse and lamprey (*Petromyzon marinus*) and one at the node represented by mouse and elephant shark (*Callorhinchus milii*). Additional lineage-specific duplications were inferred in protostomes, particularly in *Helobdella robusta*, where six duplication events were reconstructed, and in other annelid, molluscan, and ecdysozoa lineages. Thus, the GRIA-related family exhibits an evolutionary history involving diversification across bilaterians, followed by repeated lineage-specific duplications and losses (Fig. 4b; Online Resource M(b)).

#### Kainate-Type Glutamate Receptors (GRIK1-GRIK5)

Next, we examined the kainate-type ionotropic glutamate receptor family using the five human reference proteins GRIK1 (P39086), GRIK2 (Q13002), GRIK3 (Q13003), GRIK4 (Q16099), and GRIK5 (Q16478). Our orthogroup analysis recovered 78 sequences from 27 species, encompassing protostome and deuterostome bilaterians, including ecdysozoa, molluscs, annelids, nematodes, echinoderms, and chordates. The TBE and UFB phylogenies recovered several paralogue-associated groups within this dataset. Sequences assigned to GRIK4 and GRIK5 formed a strongly supported group (TBE = 1.00; UFB = 100), whereas vertebrate sequences associated with GRIK1-GRIK3 showed less consistent support at deeper branching points. In particular, a broader vertebrate-associated grouping containing GRIK1-, GRIK2-, and GRIK3-associated sequences received TBE = 0.94 but UFB = 48, indicating uncertainty in the deeper relationships among these groups. An internal vertebrate grouping containing GRIK2- and GRIK3-associated sequences, together with a GRIK1-associated sequence, received stronger support (TBE = 1.00; UFB = 100). The tree therefore supports several local relationships within the kainate receptor family, while the broader paralogue relationships remain less consistently resolved. No sequences were identified as unstable by LSI or t-index analyses, and all recovered sequences were retained for reconciliation (Fig. 4a; Online Resource J).

GeneRax reconciliation inferred 31 duplication events, 46 speciation events, and 52 speciation-loss events, no standalone-loss events. Duplication events were distributed across several bilaterian lineages. The largest concentration occurred at the ancestral node represented by Dermacentor albipictus and Argiope bruennichi, where seven duplications were inferred. Additional duplication events were reconstructed at vertebrate ancestral nodes represented by mouse and lamprey and by mouse and zebrafish (*D. rerio*), with two events at each node. Further lineage-specific duplications occurred in arthropods and other protostomes, including *Caenorhabditis elegans*, *D. melanogaster*, and *Daphnia pulex*. These results indicate repeated diversification of kainate receptor-related sequences across bilaterians, with independent expansions in protostome lineages and additional duplication events along vertebrate evolution (Fig. 4b; Online Resource M(b)).

#### Delta-Type Ionotropic Glutamate Receptors (GRID1 and GRID2)

We next investigated the delta-type ionotropic glutamate receptor family using human GRID1 (Q9ULK0) and GRID2 (O43424) as reference queries. The resulting orthogroup comprised 12 sequences from seven vertebrate species: *Anolis carolinensis, Callorhinchus milii*, *D. rerio*, *Latimeria chalumnae, Lethenteron reissneri, Monodelphis domestica,* and *Takifugu rubripes*. Both phylogenetic reconstructions recovered a strongly supported GRID1-associated group (TBE = 1.00; UFB = 100). A second group containing predominantly GRID2-associated sequences, together with a GRID1-associated sequence from Lethenteron reissneri, received lower support (TBE = 0.79; UFB = 77). The two divergent lamprey sequences assigned to GRID1 and GRID2 clustered together with TBE = 0.88 and UFB = 90. Thus, although the GRID1-associated vertebrate group was strongly supported, the relationship between the GRID1- and GRID2-associated sequences was less clearly resolved in the lamprey lineage. No sequences were identified as unstable by LSI or t-index analyses, and no taxa were removed before reconciliation (Fig. 4a; Online Resource J).

GeneRax reconciliation inferred four duplication events, seven speciation events, and six speciation-loss events, with no standalone-loss events. The root of the reconciled tree was mapped to the ancestral node represented by mouse and lamprey. Two duplication events were mapped to the ancestral node represented by mouse and elephant shark, corresponding to the gnathostome ancestral lineage. An additional duplication was reconstructed at the node shared by zebrafish and fugu (*Takifugu rubripes*), while another was assigned to the *Lethenteron reissneri* lineage. These results indicate a duplication-rich history within the sampled vertebrate GRID family, involving early vertebrate and gnathostome-associated events followed by further lineage-specific diversification (Fig. 4b; Online Resource M(b)).

#### Metabotropic Glutamate Receptors (GRM1-GRM8)

Finally, we examined the metabotropic glutamate receptor family using the eight human reference proteins GRM1 (Q13255), GRM2 (Q14416), GRM3 (Q14832), GRM4 (Q14833), GRM5 (P41594), GRM6 (O15303), GRM7 (Q14831), and GRM8 (O00222). Our orthogroup analysis recovered 92 sequences from 33 species, including anthozoan cnidarians and diverse protostome and deuterostome bilaterians. The TBE and UFB phylogenies supported several local groupings corresponding to the different metabotropic receptor subfamilies. A vertebrate-associated grouping containing GRM4-, GRM6-, GRM7-, and GRM8-associated sequences was strongly supported (TBE = 1.00; UFB = 99). Within this grouping, the four GRM4-associated sequences formed a well-supported group (TBE = 0.99; UFB = 100). The human GRM5 sequence occurred within a tetrapod-associated group containing sequences from *Anolis carolinensis, Gallus gallus*, and *Xenopus tropicalis*, which received TBE = 0.96 and UFB = 97; however, the more restricted human-chicken pairing was weakly supported (TBE = 0.52; UFB = 52). GRM2- and GRM3-associated sequences formed a broader vertebrate-rich group with support of TBE = 0.97 and UFB = 96, although one internal relationship had substantially lower support (TBE = 0.79; UFB = 37). These results support several local relationships among vertebrate-associated GRM subfamilies but indicate that some deeper paralogue relationships remain uncertain. No sequences were identified as unstable by LSI or t-index analyses, and all recovered sequences were retained for reconciliation (Fig. 4a; Online Resource J).

GeneRax reconciliation inferred 22 duplication events, 69 speciation events, and 84 speciation-loss events. Duplication events were distributed across several ancestral lineages, including the node represented by the common ancestor of Acropora muricata and Exaiptasia diaphana, where five duplications were inferred, and the node represented by mouse and lamprey, where another five were reconstructed. Three additional duplication events were mapped to the ancestral node represented by zebrafish and fugu, while further events occurred in other cnidarian and vertebrate lineages. These results indicate repeated duplication and loss-associated evolution within the metabotropic receptor family, including independent expansions in sampled cnidarian and vertebrate lineages. However, the relative evolutionary order of all eight human GRM paralogues cannot be established unequivocally from the deeper tree relationships alone (Fig. 4b; Online Resource M(b)).

#### Ionotropic NMDA Receptors (GRIN1, GRIN2A-GRIN2D, GRIN3A, and GRIN3B)

N-methyl-D-aspartate receptors (NMDARs) are ionotropic glutamate receptors that contribute to excitatory neurotransmission and synaptic plasticity. In humans, the NMDAR subunit repertoire comprises GRIN1, the four GRIN2 paralogues (GRIN2A-GRIN2D), and GRIN3A and GRIN3B. Using the seven human UniProt/Swiss-Prot reference proteins as query sequences, we investigated their evolutionary history across the 89-species eukaryotic dataset. The retained phylogenetic dataset comprised 60 sequences representing 33 species, including cnidarians and diverse protostome and deuterostome bilaterians. No identifiable homologues from non-metazoan species were recovered. The detected taxonomic distribution was not uniform across the receptor subfamilies: GRIN1-like sequences occurred in cnidarians and multiple bilaterian lineages, whereas the recovered GRIN2- and GRIN3-associated sequences were predominantly represented by vertebrate homologues.

The TBE and UFB phylogenies recovered several well-supported local groupings within the NMDAR family, although the relationships among some paralogue-associated lineages were less consistently resolved. Vertebrate-associated GRIN3A- and GRIN3B-annotated sequences formed a combined 15-sequence group supported by TBE = 1.00 and UFB = 100. Within this assemblage, a five-sequence GRIN3A-associated group was strongly supported (TBE = 1.00; UFB = 100), while a broader group containing human GRIN3B and its vertebrate homologues, together with additional GRIN3A-associated sequences, received TBE = 0.95 and UFB = 99. Thus, although the broader GRIN3-associated assemblage was strongly supported, the separation of all GRIN3A- and GRIN3B-associated sequences was not consistently recovered. For GRIN2A-associated sequences, a vertebrate group received TBE = 0.99 and UFB = 100. Similarly, a seven-sequence GRIN2B-associated group was strongly supported (TBE = 0.98; UFB = 100), although one of its internal groupings showed lower support (TBE = 0.85; UFB = 63). The human GRIN2D reference sequence was recovered within a group otherwise containing GRIN2C-associated homologues, supported by TBE = 1.00 and UFB = 100, rather than within a separately resolved GRIN2D-specific group. This placement limits the interpretation of the GRIN2C-GRIN2D relationship in the present dataset. GRIN1-like sequences displayed a broader distribution: several cnidarian-associated groups were recovered, including a four-sequence grouping supported by TBE = 1.00 and UFB = 100, while a broader five-sequence grouping received lower support (TBE = 0.88; UFB = 70). Additional GRIN1-like groupings were recovered among protostomes and chordates, including a chordate-associated group supported by TBE = 1.00 and UFB = 100 (Online Resource J). No sequences were identified as unstable by the LSI or t-index analyses; consequently, no taxa were removed before reconciliation.

GeneRax reconciliation inferred 11 duplication events, 48 speciation events, and 48 speciation-loss events, with no standalone-loss events. Duplication events were distributed across several ancestral lineages. One duplication was mapped to an anthozoan ancestral node represented by *Acropora muricata* and *Exaiptasia diaphana*, and another to the bilaterian ancestral node represented by mouse and the rotifer *Adineta vaga*. Four duplication events were assigned to the gnathostome ancestral node represented by mouse and elephant shark (*Callorhinchus milii*), followed by an additional duplication at the ancestral node represented by mouse and zebrafish (*D. rerio*). Two further duplication events were mapped to the teleost ancestral node represented by zebrafish and fugu (*Takifugu rubripes*). Additional lineage-specific duplications were inferred in Gallus gallus and *D. rerio*. Together, these results are consistent with an early origin of GRIN1-like sequences before the diversification of Bilateria, followed by substantial diversification of NMDAR-related paralogues during vertebrate evolution. However, uncertainty in deeper phylogenetic relationships and the placement of human GRIN2D among GRIN2C-associated sequences prevent an unambiguous assignment of every human NMDAR paralogue to a specific ancestral duplication event (Fig. 4b; Online Resource M(b)).

## Discussion

Across 89 sampled eukaryotic species, the human-reference-anchored analyses indicate that the glutamatergic repertoire is an evolutionary mosaic, comprising protein families with distinct taxonomic distributions and different histories of paralogue diversification. Rather than emerging simultaneously as a single functional module, the lineages represented by human glutamatergic genes combine broadly conserved metabolic enzyme families with metazoan- and bilaterian-distributed transporter and receptor families, followed by substantial paralogue diversification mapped primarily to early vertebrate, gnathostome, and bony-vertebrate ancestral nodes. Because sequence recovery was seeded exclusively with human reference proteins, these patterns specifically trace the evolutionary histories of human-reference-associated orthogroups within the sampled dataset rather than establishing the universal origins of each protein superfamily.

The human glutamate-metabolic repertoire exemplifies how ancient enzymatic machinery was partitioned and expanded across distinct cellular compartments and taxonomic lineages. In our dataset, glutamate dehydrogenases (*GLUD*), aspartate aminotransferases (*GOT*), and alanine aminotransferases (*GPT*) were recovered across multiple eukaryotic supergroups, whereas glutaminases (*GLS*) were recovered across metazoans and the ichthyosporean *Capsaspora owczarzaki*. Biochemical and physiological studies establish that these enzymes link the tricarboxylic acid cycle and amino acid homeostasis with the glutamate-glutamine cycle required for neurotransmitter replenishment [6]. However, the evolutionary trajectories of the specific human paralogues differ markedly among families. In the *GOT* family, our phylogenetic and reconciliation analyses support a pre-metazoan divergence between cytoplasmic *GOT1* and mitochondrial *GOT2* lineages, consistent with the early establishment of compartmentalized transamination in eukaryotes, such as that required for the malate-aspartate shuttle [20]. By contrast, while *GOT2* remained largely single-copy across sampled vertebrates, the divergence of human cytoplasmic *GOT1* and putative *GOT1L1* was mapped to the osteichthyan ancestor, followed by multiple secondary losses in amphibians, birds, and teleosts.

Unlike the ancient subcellular split of *GOT1* and *GOT2*, the divergences of the lineages leading to human cytoplasmic *GPT* and mitochondrial *GPT2*, as well as kidney/brain-type *GLS* and liver-type *GLS2*, were traced to early vertebrate or gnathostome ancestral nodes. This pattern is compatible with a contribution of early vertebrate whole-genome duplications (1R/2R) to the expansion of metabolic isozyme families [21], [22], but the phylogenetic and reconciliation analyses presented here do not independently establish that these duplications generated each of the inferred paralogue lineages. Nevertheless, the precise timing of the *GPT*/*GPT2* split warrants caution because of an unresolved loss inference in *Callorhinchus milii* and the presence of a third basal paralogous cluster retained in fishes and lampreys. Similarly, the deepest nodes recovered by GeneRax for *GLS* and *GPT* represent only the earliest ancestral nodes populated in this 89-species dataset and do not preclude deeper origins across unsampled taxa. Furthermore, the evolution of *GLUD1* and *GLUD2* highlights the critical dependence of reconciliation on taxonomic sampling. Although *GLUD* evolved largely as a single-copy gene across metazoans and GeneRax mapped the *GLUD1*/*GLUD2* split to the *Homo sapiens* terminal branch, *Homo sapiens* was the sole hominoid represented in our dataset. Comparative primate genomics demonstrates that intronless *GLUD2* originated via retroposition of *GLUD1* in the hominoid ancestor [15]. Thus, our tree accurately reflects the close sequence relationship between human *GLUD1* and *GLUD2* relative to murine *Glud1*, while illustrating why terminal-branch duplication assignments in sparsely sampled clades must be contextualized with lineage-specific comparative data. Alongside these vertebrate and primate events, the inference of 11 *GLS* duplications in *Bolinopsis microptera* and independent *GPT* expansions in cnidarians and arachnids indicates that metabolic gene families have undergone recurrent, lineage-specific expansions outside Vertebrata.

Whereas the metabolic enzyme orthogroups spanned non-metazoan eukaryotes, sequences in the vesicular glutamate transporter (*SLC17A6-8*) and excitatory amino acid transporter (*SLC1A1-3*, *SLC1A6-7*) orthogroups were recovered exclusively from metazoans in our search. In mammalian synapses, VGLUTs load glutamate into synaptic vesicles [23], while plasma-membrane EAATs clear extracellular glutamate to terminate signaling and prevent excitotoxicity [24]. Within our dataset, *SLC1A* homologues were detected across sponges, ctenophores, placozoans, cnidarians, and bilaterians, whereas *SLC17A* homologues were recovered across placozoans, cnidarians, and bilaterians. The absence of detected homologues in sampled non-metazoans does not prove that broader *SLC17* or *SLC1* superfamilies were biologically absent prior to Metazoa, as distant prokaryotic or microbial eukaryotic homologues may fall below human-seeded orthogroup clustering thresholds. Within Metazoa, however, GeneRax mapped the divergence separating the *SLC17A8* (VGLUT3) lineage from the *SLC17A6*/*SLC17A7* (VGLUT2/VGLUT1) lineages to the gnathostome ancestor. Similarly, the EAAT family exhibited successive duplications across metazoan, chordate, and gnathostome nodes, including the gnathostome-mapped split of the *SLC1A3*/*SLC1A6* lineage. However, pronounced discordance between TBE and UFB support at deeper *SLC1A* nodes-such as the broad grouping containing human *SLC1A1* and *SLC1A2*-prevents a definitive reconstruction of the exact duplication order that produced all five human EAAT paralogues [25].

A comparable interplay between deep evolutionary roots and vertebrate-associated diversification characterizes the ionotropic (iGluR) and metabotropic (mGluR) glutamate receptor families. The families are evolutionarily distinct, and broad phylogenetic analyses across metazoans have revealed receptor classes and lineage-specific histories not captured by vertebrate-based classifications [10]. In our human-anchored receptor orthogroups, homologues were recovered only in sampled metazoans, but their detected taxonomic breadth varied substantially among subfamilies: GRM and GRIN orthogroups were recovered across anthozoan cnidarians and bilaterians; GRIA and GRIK across protostome and deuterostome bilaterians; and GRID only in seven sampled vertebrate species. These are observed recovery patterns under our human-query-based orthogroup approach, not proof of biological absence outside those groups. This distinction matters because broader, non-human-anchored phylogenetic work reports diverse iGluR groups across metazoans [10], [26], while an earlier study placed plant glutamate-receptor-like (GLR) sequences within the iGluR clade and proposed an origin predating the plant–animal divergence[9]. Thus, the restricted distribution in our human-anchored orthogroups is consistent with limitations of query-based detection and deep sequence divergence, and should not be interpreted as demonstrating that the corresponding receptor lineages are biologically absent from unsampled or deeply diverged taxa.

Within the recovered receptor orthogroups, our phylogenetic and reconciliation analyses inferred both invertebrate duplications and vertebrate-associated diversification. Protostome-specific duplications were prominent in our GRIA and GRIK reconstructions, including six inferred duplications in *Helobdella robusta* for GRIA and seven in the arachnid ancestor for GRIK; our analyses also inferred anthozoan duplications in GRM and GRIN. Along the vertebrate stem, inferred duplications mapped to early vertebrate, gnathostome, and bony-vertebrate nodes and contributed to the expansions of the GRIA1–4, GRIK1–5, GRID1–2, GRM1–8, and GRIN1–3 groups in our dataset. These patterns are consistent with receptor-family diversification, but the cited evidence for 1R/2R genome duplication specifically supports the vertebrate GRIN2 quartet; it does not establish that all of the listed family expansions arose through those events [27]. Comparative phylogenetic analyses likewise show that glutamate-receptor evolution across metazoans is more complex than vertebrate-based classifications suggest [10].

Within the NMDAR family, GRIN1-like sequences were broadly distributed across cnidarians and bilaterians in our recovered orthogroups, whereas GRIN2- and GRIN3-associated sequences were recovered predominantly from vertebrates. This distribution is an observation from our dataset, not evidence that those subunit lineages are biologically absent from other taxa. Functionally, conventional NMDARs contain GluN1 and GluN2 subunits, and GluN3 can also assemble into NMDARs; GluN1 provides the glycine-binding site, while GluN2 binds glutamate [28], [29]. Our trees place human GRIN2D within GRIN2C-associated sequences, do not fully separate GRIN3A and GRIN3B homologues, and cluster divergent lamprey GRID1/GRID2 sequences; deeper GRIK1–3 and GRM nodes also show discordant TBE and ultrafast-bootstrap support. These are analysis-specific uncertainties. Bootstrap metrics quantify support under different resampling approaches, so discordance warrants caution about the relevant branches rather than a blanket conclusion that all local vertebrate paralogue groups are robust [18]. Accordingly, the exact branching order of these human receptor paralogues remains unresolved in the affected parts of the trees.

Several methodological considerations frame interpretation of these findings. First, because homology searches were anchored on human reference proteins, rapidly evolving or structurally divergent homologues may be less readily detected at greater evolutionary distances; the resulting taxonomic depth should therefore be interpreted as the depth recovered by this search strategy, not necessarily the full biological distribution. Second, failure to recover a homologue from a proteome may reflect incomplete assemblies, annotation errors, or orthogroup partitioning rather than true evolutionary loss [16]. Third, pruning leaves identified as unstable by the study’s LSI and t-index analyses improved topological stability in the metabolic-enzyme trees, but those pruned taxa were necessarily absent from the subsequent reconciliation input. Leaf-stability measures assess the influence of individual taxa on inferred tree structure. In the transporter and receptor trees, no leaves crossed the study’s instability thresholds, but discordance between ultrafast bootstrap and transfer-bootstrap support at deeper nodes indicates that support depends on the measure and the relevant branching patterns remain uncertain[18], [30]. GeneRax uses a species-tree-aware maximum-likelihood framework that models sequence substitutions and gene-level duplication, and loss events [19]; duplication and loss counts should therefore be interpreted in light of the supplied alignments, species tree, and inferred gene-tree relationships, rather than as certain historical events. Finally, phylogenetic placement and inferred duplication alone do not establish biochemical activity, synaptic localization, or physiological adaptation; those require functional or localization evidence.

## Conclusion

By integrating maximum-likelihood phylogenetics, tip-stability filtering, and species-tree-aware reconciliation across 89 eukaryotic species, this study reconstructs the evolutionary histories of the human-reference-associated gene families involved in glutamate metabolism, transport, and receptor signalling. Our results indicate that the human glutamatergic genetic toolkit comprises ancient, metabolic enzyme families with broad distributions across the sampled eukaryotic supergroups (*GLUD*, *GOT*, *GPT*, and opisthokont/metazoan *GLS*), metazoan-distributed transporter families (*SLC17A* and *SLC1A*), and metazoan- or bilaterian-distributed receptor subfamilies (*GRIA*, *GRIK*, *GRID*, *GRM*, and *GRIN*). Across these broad functional tiers, the specific paralogues constituting the modern human repertoire were shaped heavily by duplications mapped to early vertebrate, gnathostome, and osteichthyan ancestors, alongside a hominoid-associated retroposition in *GLUD2* and extensive, independent gene-family expansions in non-bilaterians and protostomes. At the same time, differences between TBE and UFBoot support estimates at some deeper nodes, unresolved placements of specific paralogues such as *GRIN2D*, and the inherent sensitivity constraints of human-anchored homology searches underscore that inferred duplication histories remain conditional on taxonomic sampling and phylogenetic resolution. Together, these findings provide a rigorous, critically bounded comparative framework for understanding how ancient metabolic and signaling lineages were duplicated, retained, and remodeled along the evolutionary trajectory leading to the human glutamatergic repertoire.

## Materials and Methods

This research investigates the evolutionary emergence and distribution of selected human glutamatergic reference genes, using computational methods to analyse proteins involved in glutamate synthesis, transport, catabolism/regulation, and receptor function across diverse organisms. All proteomic and protein sequence data were obtained from publicly available databases, including NCBI and UniProtKB. The UniProt FASTA sequences used as query proteins and the proteome FASTA files used to construct the local BLAST database are provided in Online Resource A. Because full FASTA headers can interfere with local BLAST database construction, sequence headers were masked and corresponding ID-mapping dictionaries were generated to recover the original sequence descriptions and annotations (Online Resource B). Because the study used publicly available protein sequence and proteome data and involved no human participants or experimental animals, ethics approval was not required.

### Species Selection

Proteome data for the selected species were collected primarily from NCBI and UniProtKB, with additional data from other public repositories. We initially assembled 107 proteomes spanning major metazoan lineages and selected non-metazoan eukaryotes for broader phylogenetic context. The purpose of this sampling was to provide broad taxonomic coverage for tracing homologous sequences corresponding to the 40 human reference protein, rather than to reconstruct the complete gene content or evolutionary history of all eukaryotic lineages.

To assess proteome completeness, we used BUSCO v6.1.1 on 107 proteomes with the Eukaryota lineage dataset (eukaryota_odb10), containing 255 single-copy orthologs [31]. The Eukaryota lineage dataset was selected because the initial sampling included both metazoan and selected non-metazoan eukaryotic proteomes used for deeper homology and phylogenetic context, allowing a consistent completeness assessment across the initial dataset. BUSCO was used as a comparative screening measure rather than as an absolute measure of genome or proteome completeness. We retained proteomes meeting the predefined completeness threshold used in the original analysis, resulting in a final dataset of 89 proteomes with BUSCO completeness scores at or above 80%. Because BUSCO scores can be influenced by lineage-specific gene content and the representation of conserved single-copy orthologs, a lower score does not necessarily indicate that the underlying genome or proteome is globally incomplete. Therefore, the BUSCO threshold was used as a quality-control criterion and not as definitive evidence of genome completeness. The number of proteomes included in each taxonomic group, together with BUSCO completeness scores, is provided in Supplementary Table S1.

### Protein selection and homology search

Human protein records were selected from UniProtKB/Swiss-Prot using established annotations and the relevant literature. Each human protein served as the initial and independent sequence anchor for its own homology search. Thus, the non-human candidate sequences analysed downstream were identified starting from a specific human reference gene rather than from an arbitrarily selected non-human sequence. The study is therefore explicitly human sequence anchored.

Human reference protein sequences used for homology searches were retrieved from UniProtKB/Swiss-Prot [17] and supplemented with curated information from the literature. The 40 focal human reference protein sequences served as the primary seed queries across the 89 proteomes. The number of initial query proteins is distinct from the number of human-associated proteins subsequently recovered and analysed. In the catabolic dataset, GOT1 and GPT were the only initial seed queries; additional human-associated family members, including GOT1L1, GOT2, and GPT2, were identified during downstream sequence recovery and included in the relevant family-level analyses. These additional proteins were not counted as independent seed queries.

We employed PSI-BLAST v2.12.0+ [32] using each of the 40 curated focal human reference proteins as an independent query. Searches were conducted for three iterations with an E-value threshold of 1e-25, and retained candidates were passed to downstream filtering (see Online Resource C).

Filtered candidates were annotated by BLASTP [33] against the UniProtKB/Swiss-Prot database [17] (see Online Resource D). A top hit was accepted for putative functional annotation when sequence coverage exceeded 80% and percent identity exceeded 40%. Functional annotation was treated as supporting evidence and not as a substitute for sequence homology, domain architecture, or phylogenetic placement [34].

BLASTP hits across all 89 species were sorted by species. Intra-species redundancy was removed using CD-HIT v4.8.1 [35] with a sequence identity threshold of 100% (-c 1.0) and default parameters (see Online Resource E). This step ensured that only unique protein sequences were retained for each species, reducing redundancy in downstream analyses.

Broccoli v1.3 [36], a phylogeny-aware orthology assignment tool, was applied to the filtered candidate sequences. Broccoli combines sequence similarity and gene-tree information to delineate orthologous groups. The resulting assignments were used to organize candidate homologs associated with each human reference gene and to distinguish potential paralogs from the focal human-associated lineage where supported by the gene tree. The Broccoli orthology assignments and resulting orthologous groups are provided in Online Resource F.

The resulting orthogroups were evaluated with InterProScan v5.73-104.0 [37] to verify conserved domains and protein-family characteristics. Automated annotation was complemented by manual inspection so that sequences lacking the expected domain architecture were excluded from the focal human-associated datasets. This step reduced the likelihood of treating broad functional similarity as evidence of homology. The InterProScan domain annotations and corresponding functional classifications are provided in Online Resource G.

For borderline or multi-domain candidates, inclusion was determined by the expected domain architecture of the focal protein family, compatibility of domain organization with the human reference protein, and phylogenetic placement within the corresponding homologous group. Sequences showing incompatible domain architectures, conflicting family assignments, or phylogenetic placement outside the focal homologous lineage were excluded from the focal dataset. Where sequence-level functional annotation conflicted with domain or phylogenetic evidence, sequence homology and phylogenetic placement were given precedence over automated functional annotation.

For downstream analysis, candidate sequences were organized according to their corresponding human reference genes and grouped into four functional categories: glutamate synthesis, catabolism, receptor signalling, and transport. Functional categories were used for biological presentation only; phylogenetic inference was performed within appropriate homologous sequence frameworks, including gene-family, orthogroup, or broader superfamily-level datasets where evolutionary relationships among the corresponding protein families had been established.

To assess receptor topology, we analysed candidate glutamate-receptor sequences for predicted transmembrane helices using Phobius v1.01 [38]. Because glutamate receptor families differ in their predicted transmembrane (TM) topology, ranging from approximately 3 TM domains in ionotropic receptors to 7 TM domains in metabotropic receptors, sequences with 2-8 predicted TM domains were retained for downstream analysis. We then used CD-HIT (-c 0.8) to reduce redundancy while preserving sequence diversity within the receptor dataset.

To explore sequence-space relationships across the four functional groups, we employed CLANS2 v2.2.2 [39], which visualizes pairwise sequence similarity using an all-vs-all BLAST-based clustering approach. Receptor, biosynthetic-enzyme, catabolic-enzyme, and transporter datasets were analysed independently. CLANS2 was run across a range of pairwise similarity P-value thresholds from 1e-15 to 1e-100. Increasing stringency was used to distinguish coherent sequence groups and to identify candidates suitable for family-specific phylogenetic analysis. Sequences that formed well-defined and stable CLANS2 clusters at the selected stringency levels were retained for downstream comparative and phylogenetic analysis. Details of the CLANS2 clustering analysis are provided in Online Resource H. The retained sets were subsequently evaluated by alignment, phylogenetic reconstruction, and reconciliation.

### Phylogenetic Tree Construction and Analysis

Following homology, domain-architecture, and CLANS2 filtering, phylogenetic datasets were assembled at the protein-family or orthogroup level, according to the evolutionary relationships being investigated. Where relationships among related human paralogues were of interest, the corresponding human reference proteins and their recovered homologues were analysed together within a family-level phylogeny. This approach was applied to the GRID, GRIA, GRIN, GRIK, and GRM receptor families. Each dataset retained the relevant human reference protein or proteins as identity anchors for interpreting the human-associated lineages. The human UniProtKB/Swiss-Prot reference sequence was retained in each dataset as an annotated reference sequence and as the identity anchor for interpreting the human-associated lineage. Functional similarity alone was not used as a criterion for inclusion. Each defined phylogenetic dataset was aligned independently using MAFFT v7.505 [40]. Post-alignment, positions with more than 70% gaps were removed using trimAl v1.5.rev0 [41] to eliminate poorly aligned and phylogenetically uninformative regions (Online Resource I).

Phylogenetic trees were constructed using IQ-TREE2 v2.2.2.7 [42], with ModelFinder [43] used to select the best-fit substitution model according to the Bayesian Information Criterion (BIC). Nodal support was evaluated with 1,000 ultrafast bootstrap replicates (UFBoot) (parameters: -m MFP -mset cpREV,Dayhoff,DCMut,FLU,HIVb,HIVw,JTT,JTTDCMut,LG,mtART,mtMAM, mtREV,mtZOA,mtMet,mtVer,mtInv,Poisson,PMB,rtREV,VT,WAG,GTR20 -mrate E,I,G,I+G,R -B 1000 --wbtl --bnni --alrt 1000 --abayes) and transfer bootstrap expectation (TBE) (parameters: -b 100 --tbe) values from 100 non-parametric bootstrap replicates [18], [44]. The corresponding IQ-TREE phylogenies are provided in Online Resource J. Maximum-likelihood gene trees were inferred as unrooted phylogenies; therefore, no specific outgroup was designated during tree construction. For gene-tree/species-tree reconciliation, the unrooted maximum-likelihood gene trees were supplied to GeneRax together with an independently derived rooted species tree [19].

Glutamate receptor sequences were analysed in separate family-level phylogenies for the delta-type (GRID1 and GRID2), AMPA-type (GRIA1–GRIA4), NMDA-type (GRIN1, GRIN2A– GRIN2D, and GRIN3A–GRIN3B), kainate-type (GRIK1–GRIK5), and metabotropic (GRM1– GRM8) receptor families. Sequences within each family were analysed together based on their homology, allowing evolutionary relationships among the human-reference-associated receptor lineages and their closely related homologues across the sampled taxa to be evaluated within a common phylogenetic framework. Potential artefacts associated with highly divergent sequences were addressed through sequential homology, domain-architecture, CLANS, and phylogenetic-stability filtering.

### Rogue taxa analysis

We employed two complementary methods, the t-index [45] and the Leaf Stability Index (LSI) [46], to identify unstable or rogue sequences that could distort phylogenetic inference. The t-index, implemented in IQ-TREE2, evaluates how frequently a taxon changes its phylogenetic position across bootstrap replicates. High t-index values indicate topological instability. We primarily relied on TBE-based t-index scores, and sequences with a t-index > 2 were classified as unstable.

The Leaf Stability Index (LSI) provides a complementary measure by estimating the frequency with which a taxon maintains a consistent phylogenetic placement across bootstrap trees. It is calculated using quartet frequencies and is less sensitive to small topological rearrangements. LSI was computed using RogueNaRok 1.0.1 with TBE-based trees [47]. Sequences with LSI scores of 0 were classified as unstable. Taxa flagged as unstable by either the t-index or LSI analysis were removed from the gene trees used as input for reconciliation to limit their potential influence on downstream inferences. The corresponding unstable sequence identifiers are provided in Online Resource K.

### Reconciliation analysis

To place the human-associated gene lineages in the context of species evolution, we performed gene-tree/species-tree reconciliation using GeneRax v2.1.3 [19] with the UndatedDL model (-r UndatedDL). Reconciliation was performed separately for each defined phylogenetic dataset so that duplication, loss, and speciation events could be evaluated without conflating distinct protein-family contexts. The primary objective was to determine the evolutionary placement of the lineage corresponding to each human reference gene.

GeneRax was supplied with the unrooted maximum-likelihood gene tree for the corresponding phylogenetic dataset and a rooted species tree derived from the Open Tree of Life [48]. The OToL topology was pruned to the taxa represented by the 89 proteomes. The rooted OToL species tree used for reconciliation is provided in Online Resource L. Using an externally curated species tree ensured that the species-tree framework was independent of the gene trees generated during candidate discovery and orthology inference. This design provides a non-circular framework for locating human reference-gene lineages within the species tree.

The substitution model used for each phylogenetic dataset was based on the best-fit model determined during gene-tree construction in IQ-TREE2 using the Bayesian Information Criterion (BIC). Reconciliation results were interpreted primarily with respect to duplication, loss, and speciation events affecting the lineage corresponding to each human reference gene.

The reconciliation output was visualized using ThirdKind v3.13.5 [49]; the TBE reconciliation visualizations and GeneRax output results are provided in Online Resource M(a) and M(b), respectively. The reconciled trees were examined to identify the deepest ancestral node at which each human-reference-associated lineage could be traced within the sampled species tree. This node was interpreted as the earliest detectable ancestral placement of the lineage in the sampled dataset, not as a calibrated divergence time.

## Statements and Declarations

### Ethics approval and consent to participate

Not applicable. This study was entirely computational and used only publicly available proteomic and protein sequence data; no human participants, animals, or identifiable private data were involved, so ethics approval and consent to participate were not required.

### Consent for publication

Both authors have read and approved the final manuscript and consent to its submission and publication.

### Authors’ contributions

MK conceptualized the work and AT carried out the research.

### Availability of data, code and materials

All proteomic and protein sequence data used in this study were obtained from publicly available databases, including NCBI and UniProtKB/Swiss-Prot. Supplementary Table S1 provides comprehensive details on the proteomic sequence data, including taxonomic classification, species or proteome counts per taxon, source, accession numbers, and BUSCO completeness scores, as well as the reference sequences used for orthogroup identification. Additional datasets, phylogenetic trees, alignments, and supplementary analyses are available at: https://doi.org/10.6084/m9.figshare.34789440.

## Acknowledgements

The authors acknowledge the Central University of Himachal Pradesh for providing computational facilities and institutional support.

